# Applying Machine Learning Technology in the Prediction of Crop Infestation with Cotton Leafworm in Greenhouse

**DOI:** 10.1101/2020.09.17.301168

**Authors:** Ahmed Tageldin, Dalia Adly, Hassan Mostafa, Haitham S Mohammed

## Abstract

The use of technology in agriculture has grown in recent years with the era of data analytics affecting every industry. The main challenge in using technology in agriculture is identification of effectiveness of big data analytics algorithms and their application methods. Pest management is one of the most important problems facing farmers. The cotton leafworm, *Spodoptera littoralis* (Boisd.) (CLW) is one of the major polyphagous key pests attacking plants includes 73 species recorded at Egypt. In the present study, several machine learning algorithms have been implemented to predict plant infestation with CLW. The moth of CLW data was weekly collected for two years in a commercial hydroponic greenhouse. Furthermore, among other features temperature and relative humidity were recorded over the total period of the study. It was proven that the XGBoost algorithm is the most effective algorithm applied in this study. Prediction accuracy of 84 % has been achieved using this algorithm. The impact of environmental features on the prediction accuracy was compared with each other to ensure a complete dataset for future results. In conclusion, the present study provided a framework for applying machine learning in the prediction of plant infestation with the CLW in the greenhouses. Based on this framework, further studies with continuous measurements are warranted to achieve greater accuracy.

## 1. Introduction

According to the Food and Agricultural Organization of the United Nations (FAO), the global population is set to reach 9.2 billion by the year 2050 (**FAO, 2017)**. To be able to feed the constantly growing population, a greater efficiency within the current farming methods is necessary. Thus, crop yield increase is one of the important targets in agriculture practices. Herbivorous insects are said to be responsible for destroying one fifth of the world’s total crop production annually **(Sallam, 1999)**. In modern-day agriculture farmers have access to sensors and mechanization that are constantly developed and conformed, enabling high automation and precision farming **(Divya and Chinnaiyan, 2019)**. These decision-support tools are directed towards more effective and efficient design and delivery of cultural practices. Precision agriculture (PA)aims to increase productivity and maximize the yields of the crops**(Keswani et al., 2019)**.Integrated crop management is used to enhance crop productivity and farm income through crop management, nutrient management, and pest management. Applying machine learning (ML) to those sensor data and management systems turning it into artificial intelligence-enabled programs providing recommendations and insights on the spot.

A lot of techniques were used to understand the rules and relationships from diverse data sets, to simplify the process of acquiring knowledge from empirical data. These techniques perform well on artificial test data sets, the main goal is to make sense of real-world data **(Khattab et al., 2016)**. Machine learning (ML) offers an alternative to the conventional engineering flow when the problem is too complex to develop a solution with guarantees. On one hand the approach has the disadvantages of producing black-box-solutions that are not interpretable, so they are only applicable to a limited set of problems **(Brynjolfsson and Mitchell, 2017; Simeone, 2018)**.Several research works describe different automation practices like wireless communications, IOT, ML, AI and Deep Learning. **Jha et al**.**(2019)**describe the application of a range of ML algorithms to problems in agriculture, with different problem sets and outcomes.**Chtioui et al**.**(1999)**used a generalized regression neural network (GRNN) as well as linear regression (LR), also referred to as multiple linear regression (MLR). Their applicability for leaf wetness prediction and forecasting several plant infestations were measured.

This harmful pest is one of the major key pests that attack multiple crops in Egypt around the year **(Amin and Salam, 2003)** and known to develop resistance towards common chemical insecticides **(Abou-Taleb, 2010)**. The aim of the present work lies specifically in targeting the Egyptian cotton leafworm (CLW, *S. littoralis*). Applying ML to predict the crop infestation with this insect may provide a powerful tool to control the insect in a timely and effective way to prevent its damaging effects on the crops.ML implementation can be broken down into two major parts; datasets and algorithms. In the present work, dataset is provided by sensors distributed in the greenhouse (temperature, humidity). The algorithm for the prediction of the insect behavior will be constructed with the focus on regression analysis and decision tree analysis with linear and non-linear kernels. The output of this prediction algorithm is planned to be eventually connected to an automatic system for the pest control.

## 2. Methods

### 2.1 The greenhouse

The experiment was carried out at the Nabat Farms in Al Mansouryah, Giza, Egypt, hydroponic greenhouse. Crisphead lettuce (*Lactuca sativa* var. capitata L. nidus jäggeri Helm, common name: Batavia) is cultivated periodically so all plant’s ages are always present in the hydroponic greenhouse. Eight light traps were fixed in the hydroponic greenhouse. The traps placed above the plants with about one-meterhigh and were kept on this level until the end of the experiment. The light traps used to catch the moth of CLW. The population density of the CLW was estimated weekly by collected the moth from the light traps, counted and recorded. Daily data of minimum, maximum temperatures and relative humidity were obtained from air temperature and humidity sensors hanging in the hydroponic greenhouse and connected to WeatherLinkapplication.

### 2.2 Dataset and features

The dataset concludes 130 records between 09/2017 and 02/2020 with each record representing one week of measurement. The plantation works as a continuous process with new plantations and harvests every day. It takes 35 to 40 days for each individual plant from seedling to harvest. This leads to a constant state during the entire time of measurement where there are always young and old plants living simultaneously in the greenhouse as well as other plants that will not be analyzed during this research. Table 1 shows the original features of the given dataset, which were later used to derive additional features as can be seen in Table 2.

**Table 1.**
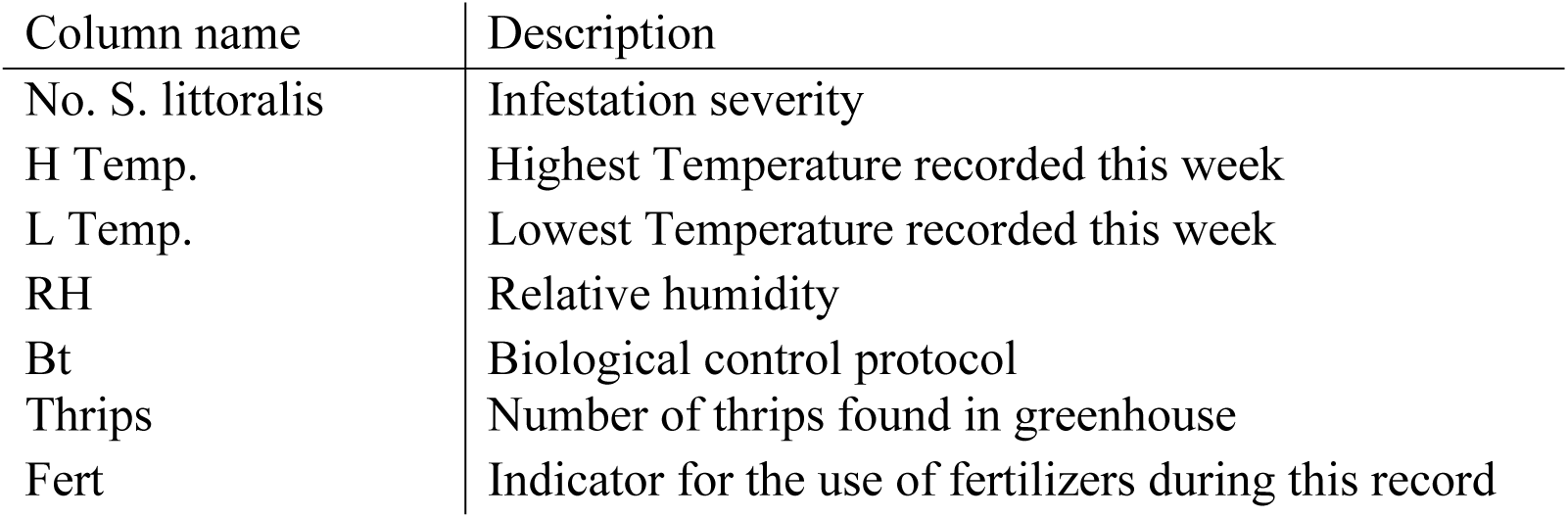
Original features for cotton leaf worm dataset

| Column name | Description |
| --- | --- |
| No. *S. littoralis* | Infestation severity |
| H Temp. | Highest Temperature recorded this week |
| L Temp. | Lowest Temperature recorded this week |
| RH | Relative humidity |
| Bt | Biological control protocol |
| Thrips | Number of thrips found in greenhouse |
| Fert | Indicator for the use of fertilizers during this record |

**Table 2.** Additional derived features for cotton leaf worm dataset

| Column name | Description |
| --- | --- |
| Tmean | Weekly average temperature |
| GDD | Growing day degree |
| lastBt | Bt on previous week |
| lastIS | Infestation severity on previous day |
| meanlast3IS | Average of infestation severity of last three weeks |
| diffTHigh | Fluctuation in highest temperatures in the |
| diffTLow | Fluctuation in lowest temperatures in the |

The original features comprise the infestation severity (IS) which representing by the total number of CLW moth captures over the period of one week for the entire period of the study. Additionally, the highest, lowest temperature and the relative humidity were recorded. Due to the commercial nature of the greenhouse in which the study was executed, a biological control agent *Bacillus thuringiensis* var. kurstaki (Bt) was applied to control the pest and included in the study. As insects could affect the activity and behavior of other co-existing insects, the column “Thrips” shows the number of *Frankliniella occidentalis* Perg, the other insect captured during the same period. The column “Fert’ is as an indicator to know when fertilizers were applied in the greenhouse to measure the effects of these fertilizers on IS.

The derived features were calculated from the original features and were used to improve the accuracy of the implemented algorithms. The derived temperature fluctuation calculations different high temperatures (diffTHigh) and different low temperatures (diffTLow) in Table 2 are necessary as each record represents a whole week, therefore allowing only one temperature measurement per week. This feature is introduced to the dataset to represent the fluctuation in the temperature during the measurement period and tune the ML algorithm. For these features, the temperature data of the entire period was collected. The highest and lowest inevery week were compared and the difference between the highest THigh per day and the lowest THigh per day was recorded in the fluctuation feature. The same method was used for the lowest temperatures.

The calculated Growing Day Degree (GDD) feature was used in previous literature **(Hannukkala, 2007)**. The GDD is calculated as GDD = T_mean_-10 and serves in the description of environmentally favorable conditions for plant infestation. As the greenhouse is operated continuously without planting seasons, the Accumulated GDD is not used as the accumulation cannot be reset. The previous infestation severity is difficult to track, as the measurements were taken weeklyand the previous record thereby represents the last week. As the biological control protocol can be activated during that time, the feature importance could be affected. Nevertheless, they will be used in the analysis.

### 2.3 Machine learning

To apply any sort of machine learning methods a dataset needs to be retrieved where each row of data represents an observation about something in the world. For all models, the dataset is divided into a train and a test set, both consisting of the features of analysis and the Key Performance Indicator (KPI). The train set is used to fit the model and define the relationship between the input features and the KPI, whereas the test set is used to measure how accurate the model is predicting the output given the test features. Different types of algorithms and models can help achieve different goals, while in their core they are all ways of figuring out what drives the changes in the (KPI) of the application.

An overview of the ML process can be seen in fig.1. It starts with the data collection and feature extraction to prepare the dataset for analysis and derive additional calculated features. Every combination of ML and algorithm and feature set was used for fitting a model. For each resulting models the predicted values were derived and evaluated in comparison with the other results. For the ML algorithms, it is distinguished between classification and regression algorithms **(Pentakalos, 2019)**.Classification is about predicting a label. It is the method of approximating a mapping function from input to discrete output. As this study aims to predict the quantity of the infestation over time, a classification approach is not taken. Regression is about predicting a quantity. It is the method of approximating a mapping function from input to a continuous output, such as an integer or floating-point value. These are often quantities, such as amounts and sizes. This is more appropriate to the present study.

**Fig.1.**
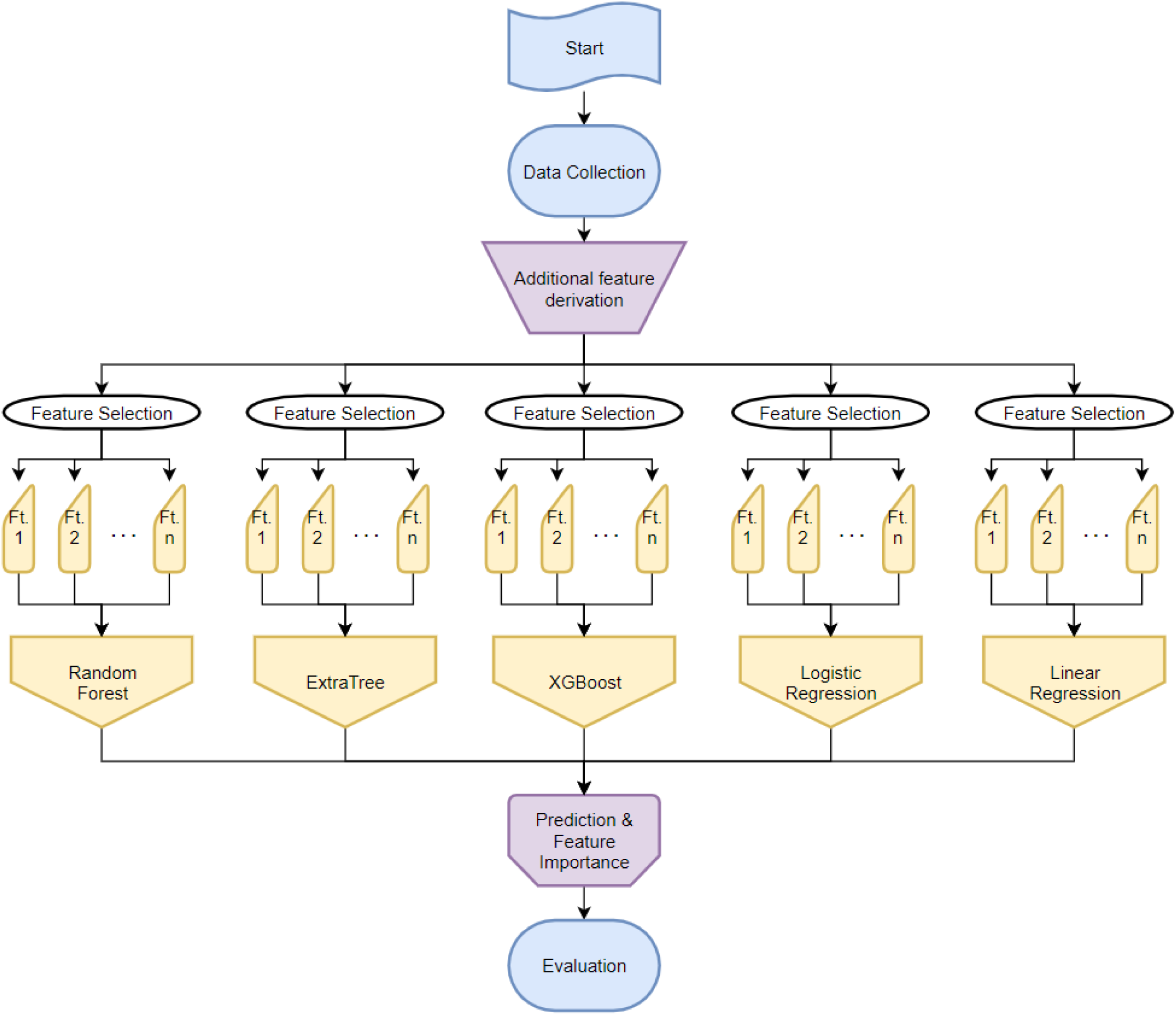
Machine learning process - Flowchart

To rank the performance of the ML models the error functions for regression algorithms are introduced. These error functions typically compare the predicted values with actual data points provided from the test set. The most popular error functions for these types of work are the Relative Root Mean Square Error (RRMSE) and the mean absolute percentage error (MAPE) **(Liakoset al., 2018)**.RRMSE is the relative standard deviation of the residuals. These prediction errors measure how distant from the actual values the predicted values are. It is also a measure for how spread out they are. It simply explains how concentrated the data points are around the line of best fit and is also commonly used in climatology, forecasting, and regression analysis to verify experimental results. With d_i_ being the actual value and f_i_ being the forecast value, the formula is given by:

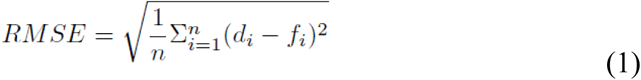

The RMSE is then divided by the average of the actual values to get the RRMSE for analysis. MAPE is a measure for how accurate a forecast system is. With d_i_ being the actual value and f_i_ being the forecast value, the formula is given by:

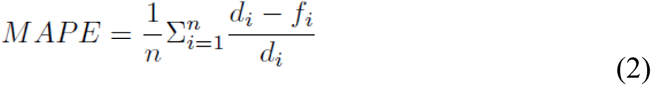

Five different algorithms were implemented and the RRMSE and the MAPE were recorded for the analysis. The SVM was not tested as it proved to be inefficient for these use cases. For each ML method five different feature combinations were captured and used for this analysis.

- In the first approach of this analysis, all the observed features from Table 1 were used and the model was enhanced by the GDD features. Information on previous IS and the temperature fluctuations was not introduced at this point.
- For the second approach, the data of the previous IS as well as the averaged IS over the last three weeks was introduced into the dataset.
- A similar behavior was recorded in the third approach, when the temperature relating data was introduced.
- For the fourth approach, the information about the thrips was excluded from the model to test the indirect impact of these insects on the cotton leaf worm.
- Lastly all features were selected at once and the predictions were determined.

## 3. Results

### 3.1 Variation in some features with time

The changes of some features with time were depicted in Figs 2, 3, and 4. The IS over time in each season is shown Fig 2. The IS as shown was not linearly increasing. This is due to the climatic conditions of the greenhouse and the biological control protocol (Bt) applied. The changes in relative humidity as well as the highest and lowest temperature for the study years were depicted in Figs 3 and 4. The highest and lowest temperature of the day is fluctuating throughout the weeks. This plot was created with the weekly recordings from the dataset. Peaks in the IS can be observed in 2018 as the control protocol was not yet tested effectively. It is clearly visible that the protocol had a more positive effect on the plants in 2019. It is notable that the relative humidity in 2018 was much higher than in 2019 which could also have led to the increased IS in 2018.

**Fig2.**
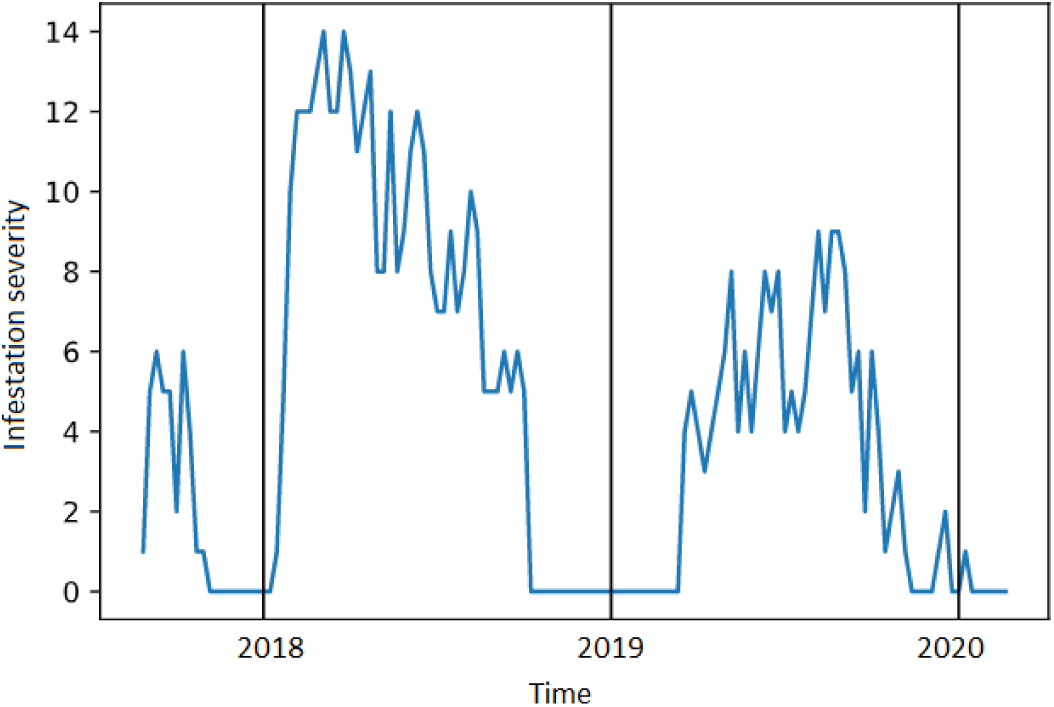
The infestation severity (IS) of CLWchanges with time

**Fig.3.**
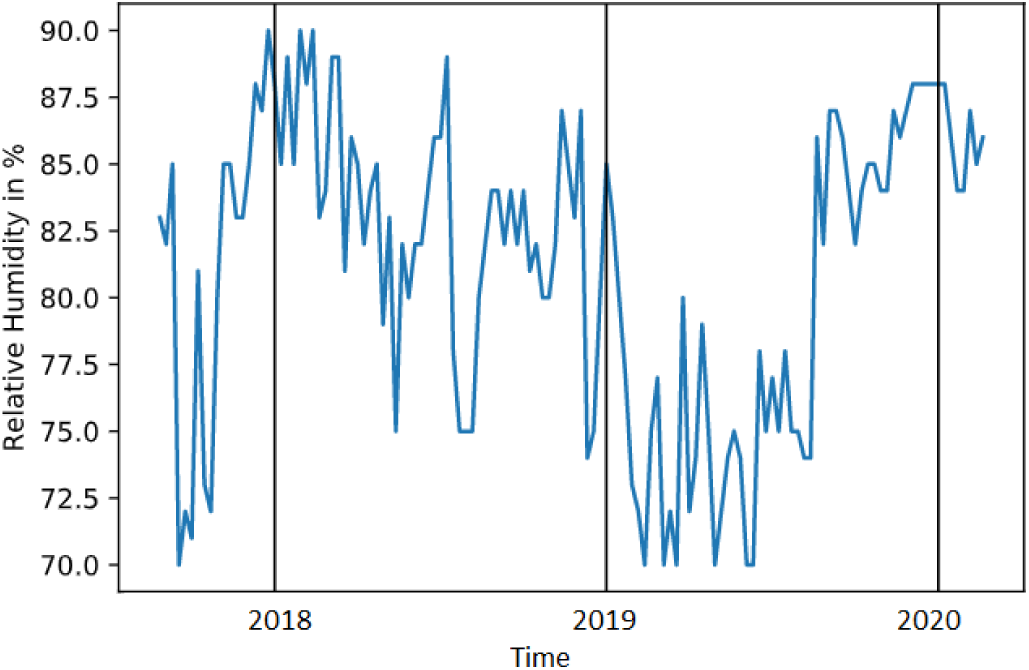
The relative humidity changes with time

**Fig.4.**
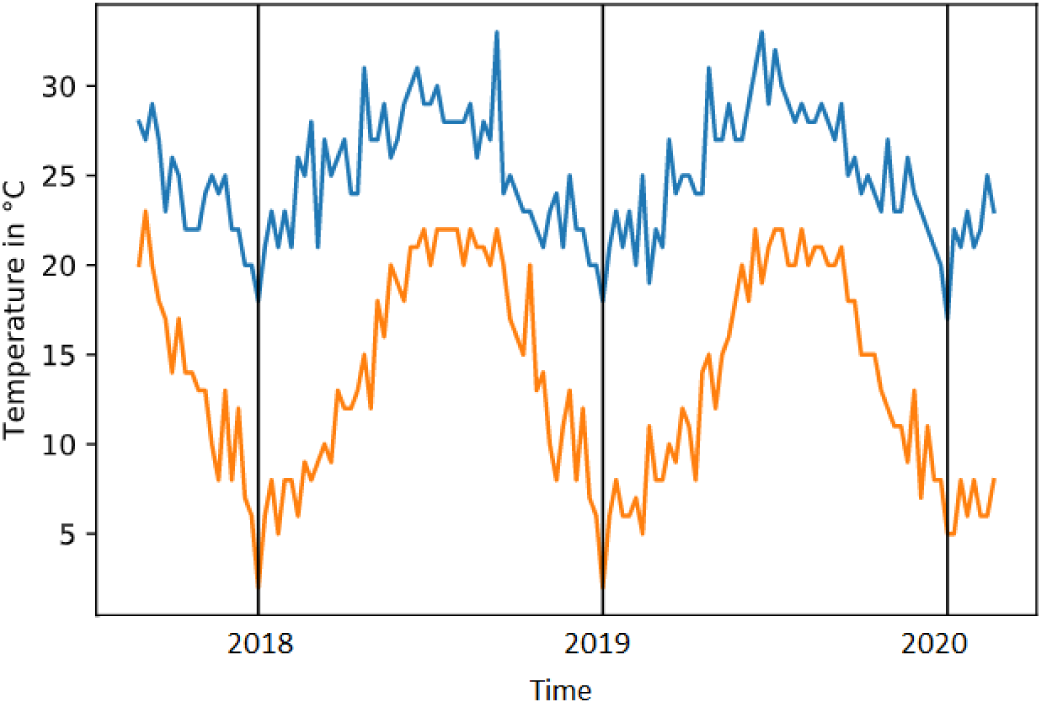
The highest (upper trace) and lowest temperature (lower trace)changes with time

### 3.2 Data Analysis

The dataset was split into training and a test sets. In the following analysis, the split is 80:20 as is typical in most ML analysis. Table 3 shows the results obtained for the predictions of the IS using each algorithm. In the first part of this analysis, all the observed features from Table 1 were used and the model was enriched by the GDD features. Information on previous IS and the temperature fluctuations was not introduced at this point.

**Table 3.**
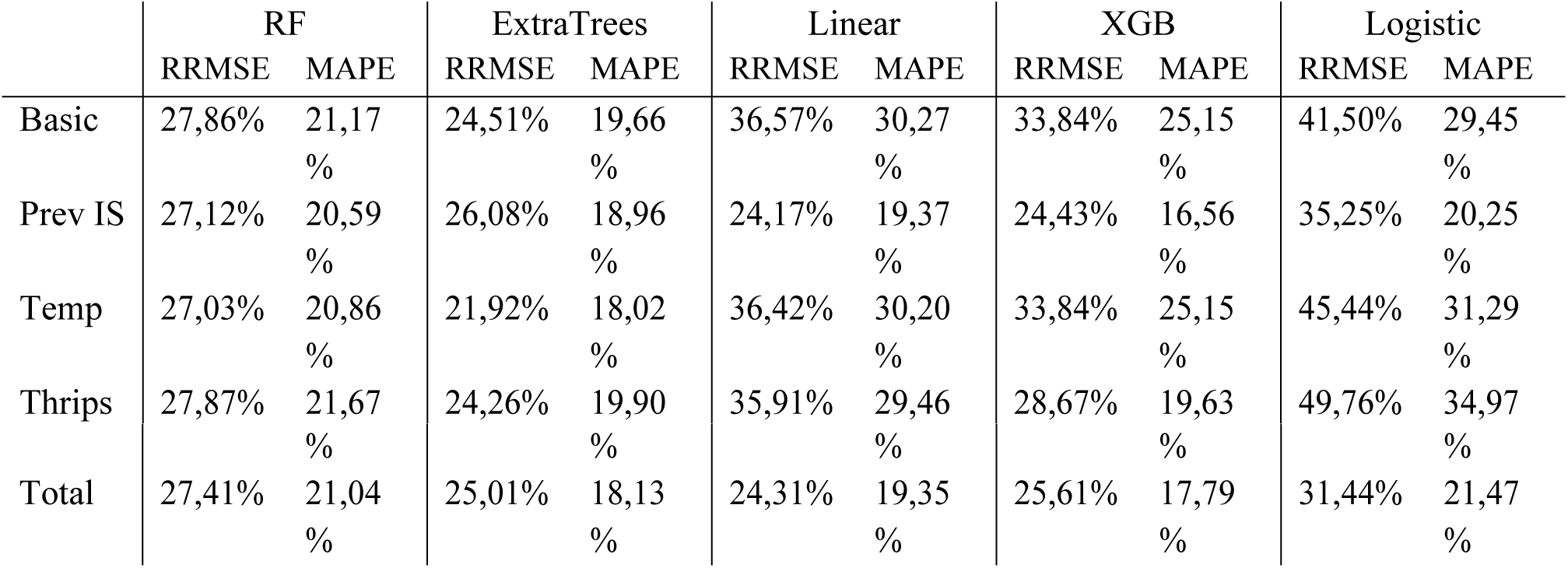
ML analysis on cotton leaf worm with random sampling

First the baseline was established by calculating the predicted error with the original observed features from Table 1. This resulted in errors between 19.66% (Extra Tree Regressor) and 30.27% (Linear Regression). The next subsections analyze the effect of each feature on the model to determine the best combination of features to achieve optimum results. Random sampling was chosen in all approaches as the Bt protocol was adapted with time and the records are all equal in terms of plant age and count. The last row of the table shows the results obtained when all features were combined. This includes the previous IS, the number of thrips as well as the temperature fluctuation and leads to stable results throughout the model.

For the second approach, the data of the previous IS as well as the averaged IS over the last three weeks were introduced into the dataset. This inclusion led to slight improvements in the prediction model. A similar behavior was recorded when the temperature relating data was introduced in the third approach. The fluctuation of the temperatures within each week seems to have a minor effect on the prediction algorithm. Lastly the information about the thrips was excluded from the model to test the indirect impact of these insects on the CLW. This step resulted in a better performance of the XGB and poorer performance in the Logistic Regression leaving the other three algorithm performances unchanged.

In the second approach, information about the previous IS was introduced to the dataset before fitting the model. This had a positive effect on all models, especially on the regression algorithms and the XGB. Adding this feature especially improved the regression algorithms as they assume a linear relationship between the inputs and the number of CLW, thus by adding usual features, they are assigned a strong weigh. The fitting model becomes more accurate. As such a useless feature has a direct negative impact on the model. The XGB improved far more than the other decision tree algorithms as boosting reduces variance and bias by using multiple models (bagging). Thereby, it trains the subsequent model by telling it what errors the previous models made using the difference between the predicted and actual values. Given the weakness of the base learner, the errors of previous models are used for the subsequent models to build upon. As a result, this feature should be introduced in the final model to ensure better results.

The third row of Table 3 shows the effect of introducing calculated temperate features to the model. This includes the temperature fluctuation features as well as the mean temperatures of the past weeks. The average temperature of the past records is not significant in the described information from three weeks ago and the insects only take days to hatch. However, these features only have a slight impact on the results. Only, in the case of the RF and the ExtraTree, they lead to a slight improvement, possibly by reducing the variance of the model. The feature is introduced to the model as well but is not expected to have a high feature importance. Increasing the records by measuring every day makes this feature obsolete and gives a better understanding about the impact of the temperature on the CLW.

The impact of the captured thrips on the base observation in the greenhouse is analyzed, by removing this feature from the basic model in the fourth row of Table 3. Removing this additional feature from the model has almost no impact on the model, except for an improvement of the XGB. The feature is tested again in the next subsection in combination with the other features.

The impact of the biological control protocol on the base observation in the greenhouse is analyzed, by removing this feature from the basic model. As this feature is essential for describing the current state of the greenhouse, removing it would have a strong negative impact. The Bt. is a reaction to the CLW affecting the plants and will therefore be difficult to integrate in a long-term prediction model. However, for 2 – 4 weeks of prediction durations it should always be included, as goes for other environment related features.

The last row in Table 3 shows the results when all features were combined, as they improve the model. As these results appear slightly less performing than the second row, additional tests were conducted by now removing the thrips and the temperature features while leaving the other as well as the previous IS. None of these combinations performed better than the second row, indicating that the model is not as stable as anticipated with the given features. The optimum stable result achieved for this dataset is a combination of all features with prediction errors ranging from 17.79% to 21.47%.

The prediction error is captured in two ways, the Relative Route Mean Squared Error (RRMSE) and the Mean Absolute Percentage Error (MAPE). In this experiment the RRMSE is higher than the MAPE by at least 5%. By squaring before averaging the errors in the RRMSE more weight is put on the records with larger errors. For this dataset the difference is up to 5-15% (e.g. Logistic Regression baseline, MAPE 29.45% and RRMSE 41.50%). This indicates that the predictions in the models have multiple outliers. Figs 5, 6, 7 shows the prediction result when using Logistic regression on the dataset with all features included, where Fig. 5 and Fig. 6 are the absolute error and the squared absolute error of the prediction. When averaging the errors from Fig. 7 it is expected that the errors at record 59 and 103 have a higher impact on the overall error, leading to an RRMSE of 31, 44% instead of 21,47% with the MAPE. The outliers have a much bigger impact on the overall error calculation.

**Fig. 5.**
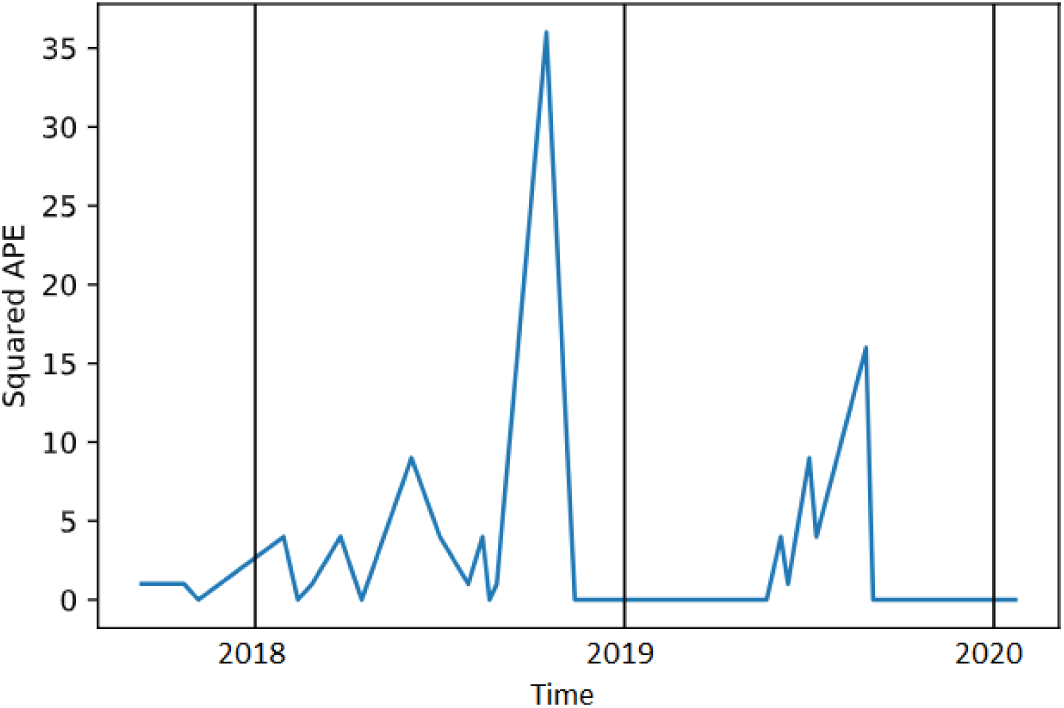
The squared absolute error

**Fig. 6.**
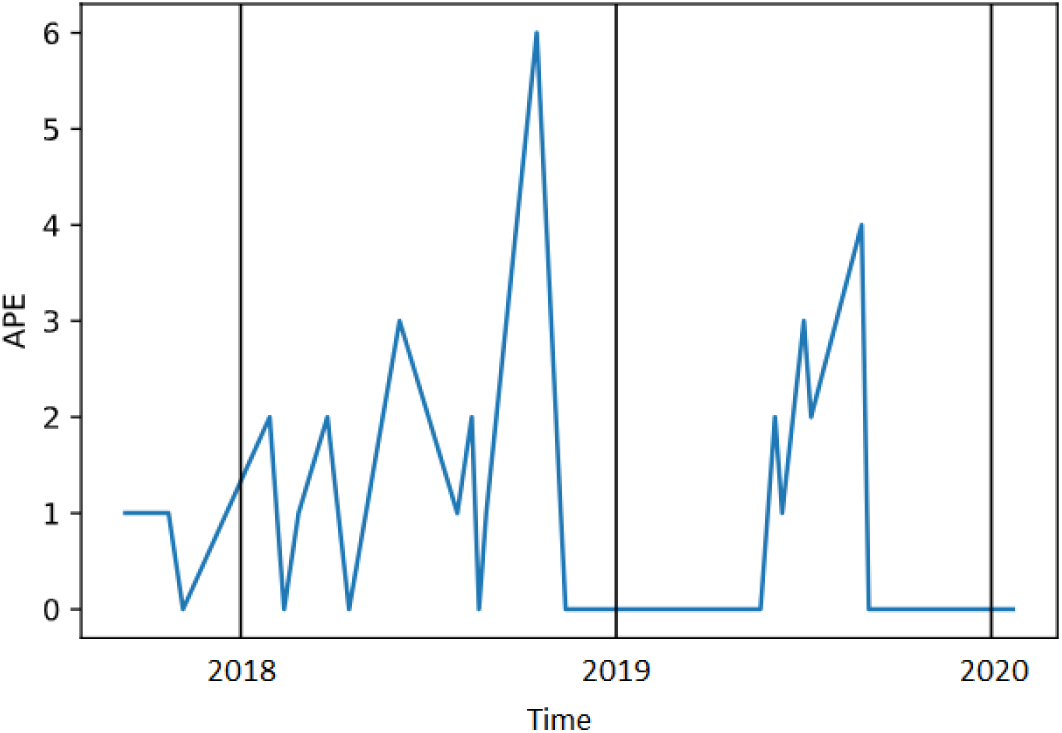
The absolute errors

**Fig. 7.**
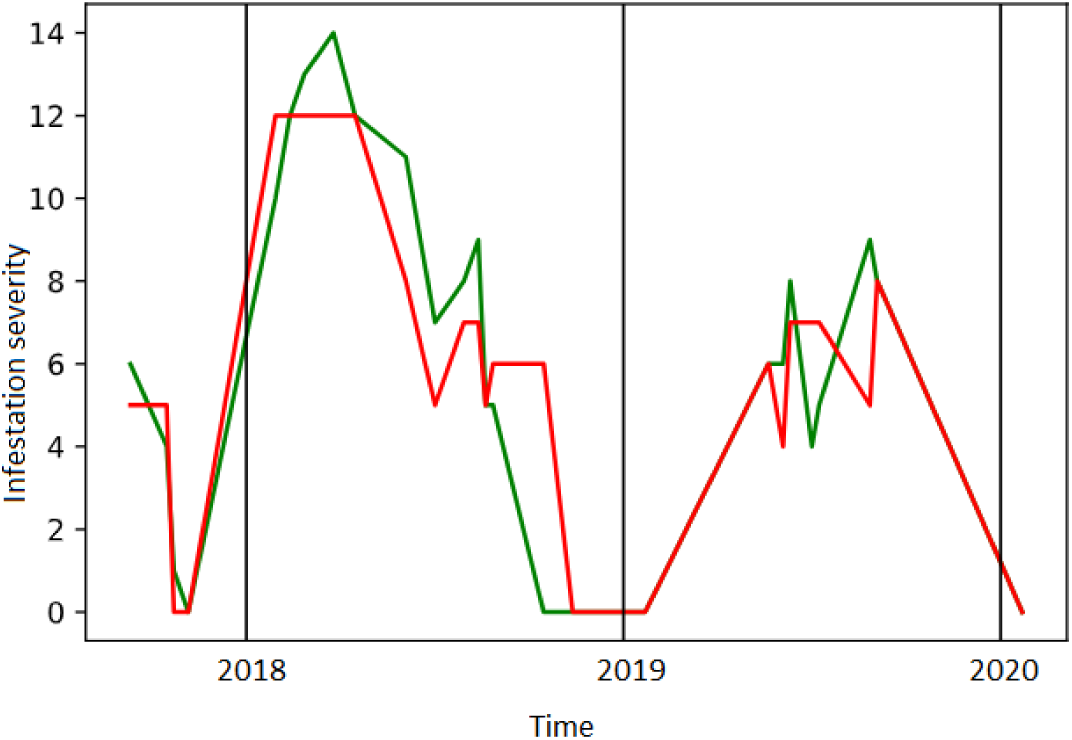
The output of ML predictions of IS (red) and the acutal data (green)

The feature importance distribution of the different algorithms is shown in Fig.8, where each line represents the decision tree algorithm and the feature importance is plotted in %. Looking at feature importance validates the above assumptions. The temperature features do not appear to have been relied on strongly in the decision tree algorithm which is why this feature did not impact the overall model strongly. Looking at the previous IS features it is also clear that the models rely strongly on these features therefore leading to improvements of the model, however the XGB relies less on them than the other two algorithms. As the XGB performs best in these cases, its importance structure appears to be more adequate for this dataset. The XGB gives higher importance to the Bt, while also considering the other features. At the same time, the other models give a very high importance only to some features while disregarding the rest which could skew the model.

**Fig. 8.**
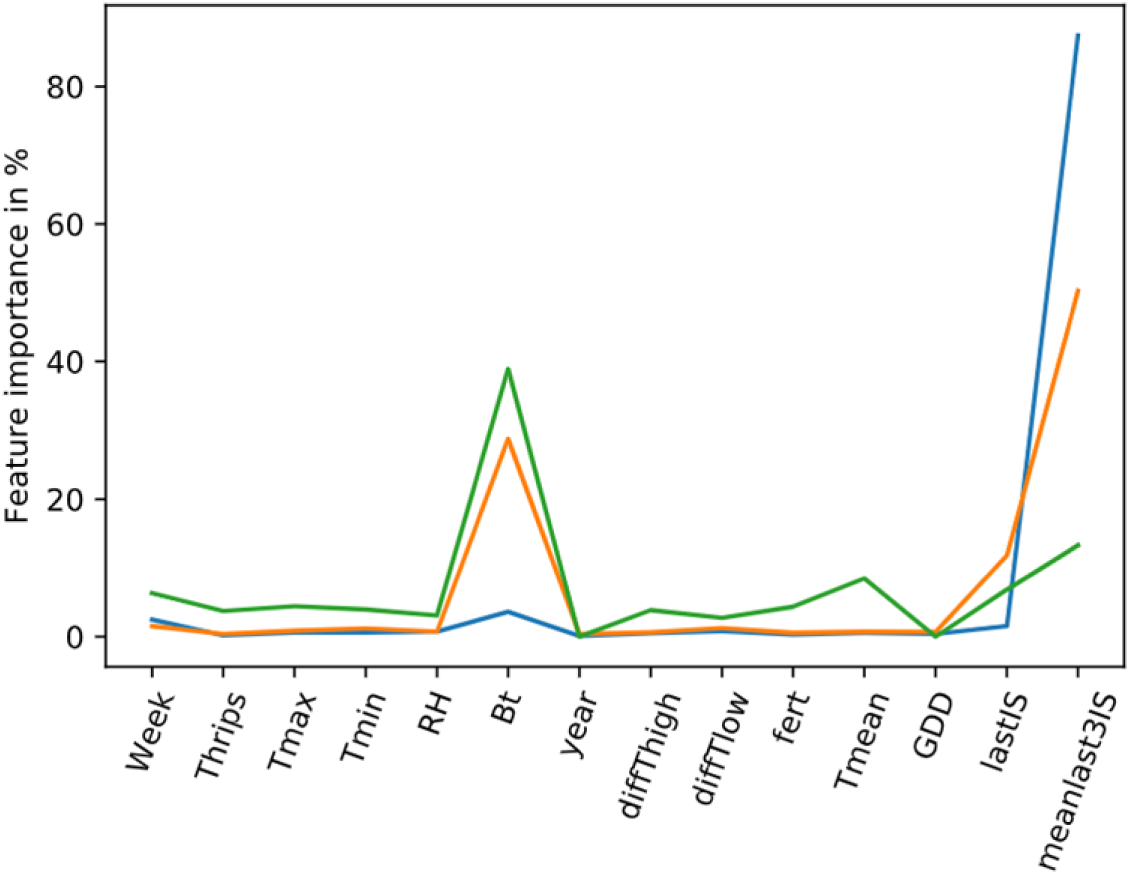
Feature analysis (importance) with decision tree algorithms

## 4. Discussion

The agriculture sector faces multiple challenges linked to infestation and pests as well as improper soil treatment and water systems and many others. Research is being conducted to address these issues. Artificial intelligence with its vast learning capabilities has become a major tool in the race to solve the agriculture related problems **(Bannerjee et al., 2018)**.Precision agriculture (PA), is the scientific field that uses data intense approaches to drive agricultural productivity regarding the environmental impact and the chemicals used **(Konstantinos et al., 2018)**. It is the application of technologies and principles managing multiple aspects of agricultural production to improve crop performance and quality **(PierceandNowak, 1999)**. Reducing the number of chemicals used in plant protection products, ultimately reduces the levels of residuals found on our food **(De Baerdemaeker, 2013)**.

Building a model for prediction of plant infestation with pests is of paramount importance in control these pests either in greenhouses (**Chiu et al., 2019**) or fields (**Skawsang et al., 2019**). Such a model may provide a tool for the perfect timing in the prevention of the damaging effects of the pests on the plant which leads to an increase in the crop yield. Moreover, it could minimize the efforts exerted to control the pests which led to a reduction in the costs. In the present study, one of the most damaging pests for several crops; CLW has been studied during the period of cultivation in one of the commercially available greenhouses. Five different ML algorithms have been implemented using the insect and the environmental original and derived features to predict crop infestation with this pest. The comparison between the output of the different algorithms were presented and the feature importance for further analysis was showed. The classification algorithms SVM/SVR are not considered, as it is normally used in binary classification problems, where the probability of an outcome occurring is the matter of interest. This method cannot be applied for this application as the dataset does not contain multiple recurring information with slight variations as is the case with classification problems. Regression algorithms are more suitable for predicting non-discrete outputs as these can be any real number, range from negative infinity to infinity given that the regression line is a continuous function.

It was showed that, the regression algorithms perform less than the decision tree algorithms, with the Linear Regression performs slightly better than the Logistic Regression. As the weights of the features are important in this use case, the decision tree algorithms achieve more promising results. This could be due to ability of decision trees to weigh certain features as more important than others, rather than assuming that features have linear relationships to the IS. This is especially the case when information on previous IS were not provided in the model and resulted in MAPEs of 29.45-34.97%. These features correlate strongly to the current IS and the lack of this information leaves the model with a strong gap. The Decision Tree algorithm was able to tackle this lack of information by increasing the importance of the other features. While the RF and the ExtraTree deliver relatively stable results, the errors of the XGB fluctuate with the number of features introduced as it does not yet deliver the expected optimum tradeoff between variance and bias **(Hastie et al., 2009; SuchithraandPai, 2019**). At the same time the other decision tree algorithms lack enough records to form a deep enough decision tree and achieve results of higher accuracy.

Given the insights from Figs2 - 4 and the previous analysis, the infestation period is expected to behave similarly in the years 2020 and 2021 and ream peaks in the warmer months from May to August. With an optimized Bt schedule, the peaks are expected to decrease significantly as can be seen when comparing 2018 and 2019.Compared to previous research in the field precision agriculture, this model shows a relatively better performance with a lot of potential for improvement as error rates of as low as 1-2% can be achieved with additional measurements. Especially given that it was possible to achieve error rates of 16% - 17% using additional derived features (e.g. the new derived features on previous IS) than the standard approach using only observed features in the ML models. To ensure more validity of the approach used in this research, the system is planned to be applied on multiple crops measuring different pests simultaneously.

## 5. Conclusion

Based on the discussion above, the dataset is best described by a decision tree algorithm. The XGB delivers more reliable and stable results in terms of the distribution of the feature importance. The additional derived features had a positive effect on the predictions as they did not lead to overfitting the model. It can be assumed that the future expansion of the project is beneficial for farmers. Thus, Pesticides or other methods of pest control can be utilized in more targeted manners, which leads to a reduction of the costs of the crop and an increase of the yield produced.

## Acknowledgement

Great thanks are due to the officials and workers in the Nabat Farms in Al Maansouryah, Giza, Egypt, for offering the opportunity to carry out the present assay in their greenhouse. This work is funded by the Information Technology Industry Development Agency (ITIDA), Information Technology Academia Collaboration (ITAC) Program, Egypt – Grant Number (CFP# 163)”

